# Exploring B and T-cell epitopes for constructing a Novel Multiepitope vaccine to combat emerging Monkeypox infection: A Reverse Vaccinology approach

**DOI:** 10.1101/2022.12.09.519581

**Authors:** Hassan Yousaf, Anam Naz

## Abstract

**Background:** While the whole mankind is resurrecting from the recent Covid-19 pandemic, new cases of the monkeypox virus have been reported inflicting serious threats to the public health. Monkeypox is a newly emerging, zoonotic orthopoxvirus having similar symptoms as that of the smallpox. So far, no approved treatment and therapeutics are in line to fight the infection.

**Methodology:** Therefore, in the present study, we have deployed a computational pipeline. We have retrieved the helper T-cell lymphocytes, cytotoxic T-cell and B-cell inducing epitopes by targeting the cell surface binding protein of the virus and further filtered the high-quality peptides based on their immunogenicity, antigenicity and allergenicity. After subsequent steps, we constructed and validated the tertiary structure of vaccine and analyzed its molecular interactions with toll like receptor-2 (TLR-2) and toll like receptor-4 (TLR-4) through molecular docking and the atomic movements and stability through molecular dynamics simulation approach. Moreover, C-IMMSIM server was used to evaluate the immune response triggering capacity of the chimeric vaccine through the immunoglobin profile.

**Conclusion:** The conducted *in silico* study concludes that the surface protein of monkeypox virus is one of the major culprit antigens in mediating the disease. Hence, our study will aid in the better formulation of vaccines in future by targeting the suitable drug or vaccine candidates.

## Introduction

Monkeypox virus (MPXV) is the aetiological agent of the zoonotic infection known as monkeypox. The virus was first discovered in 1958 while experimenting with monkeys at the Statens Serum Institute in Copenhagen, Denmark, hence they have been reported as natural hosts of the virus. However, a few other natural hosts include: rope squirrels, tree squirrels, Gambian pouched rats, and dormice [1]. Since the first isolation of monkeypox virus, its very first case was reported in an infant of nine-months old during the period of 1970 in the DRC (Democratic Republic of Congo) but the infection was primarily contained in the African region with merely a total of 48 cases being reported in the forthcoming decade for six African regions (1970-1979) [2]. But the virus re-emerged with a rather massive public health threat when the first few cases were reported in Canada=13, Spain=7, and Portugal=14 on May 18^th^ 2022. Since then, the cases were reported in different countries including Australia, Germany, Netherlands and United Kingdom [3, 4]. Monkeypox virus is a DNA orthopoxvirus from the family of Poxviridae having a large size of 200-500nm. The virus is brick-shaped having a lipoprotein envelope, and a linear double-stranded DNA genome (figure 1).

**Figure 1.**
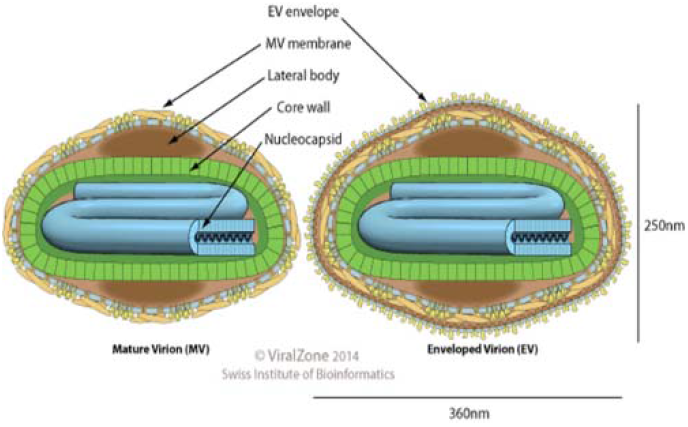
Structural presentation of Monkeypox virus. [Courtesy: ViralZone]

As of now, the virus is classified into two clades: Central African Congo Basin clade and West African clade, whereas the former has been more prevalent with a high number of human-to-human transmission cases being reported so far [5, 6]. The MPXV infection is zoonotic and primarily spreads through animal-human contact, human-human contact through close interactions including the bodily fluids, infected objects like bedding, respiratory droplets, sexual contact and contact with wounds and lesions [7]. People with monkeypox normally get a rash which initially looks like pimple or an itchy blister and these are usually located near the genitals (penis, testicles, vagina), anus and/or sometimes on hands, mouth and chest. Other symptoms include severe headaches, fever, fatigue and respiratory issues such as nasal congestion and dry cough (CDC Report). The virus can be diagnosed by real-time polymerase chain reaction of the samples including dry swabs of lesions and ulcers. Seven endemic nations have reported 1408 suspected and 44 confirmed cases between January and June 2022, resulting in 66 fatalities. Cameroon, the Central African Republic, the Democratic Republic of the Congo, Gabon, and Ghana are among the nations where monkeypox is endemic (identified in animals only). The scenario is changing, and WHO predicts that additional cases of monkeypox will be detected as the outbreak advances and surveillance increases in both endemic and non-endemic countries [8–11]. Although there is no approved vaccine or drug therapy for the disease by FDA yet but recently JYNNEOS; a safe profile vaccine was recommended by the Advisory Committee on Immunization Practices (ACIP). Though it should be kept in mind that this vaccine was recommended as a safe alternative to no treatment of monkeypox and hence it is not composed of the monkepox virus itself and incase of mutations in the virus, the vaccine will lose its effectiveness. Albeit, conventional vaccines have proven to elicit potent neutralizing antibodies and a robust immune response to infectious diseases; but they are extremely costly, time-consuming and presents with a number of side effects. Thus, in this study, we have proposed a peptide-based chimeric vaccine using the immunoinformatics approach.

## Materials and Methods

### Retrieval of Protein Sequence and Preliminary Analysis

The full-length amino acid sequence of the cell surface-binding protein of monkeypox virus (strain Zaire-96-I-16) was retrieved from the UniProt database under the entry of: **Q8V4Y0 · CAHH_MONPZ** in FASTA format. For validation of protein antigenicity, we deployed the *VaxiJen* v2.0 server [12] with a default threshold of 0.4.

### Prediction of Cytotoxic T-cell (CTL) and Helper T-cell (HTL) Epitopes

The antigenic protein was fed into the IEDB server, where we employed the Tepitool [13] in order to retrieve the MHC-I (CTL) binding peptides by selecting a panel of 27 most frequent alleles. The prediction was based upon the percentile rank in which the peptide is ranked by comparison of predicted binding affinity of epitopes with a panel of random epitopes from SwissProt. Hence, a lower percentile rank indicates better binding of the peptide with respective alleles. Additionally, we modified the settings to retrieve low number of peptides (9mers). While the output of the predictions is quantitative, there are systematic deviations from experimental IC50 values. The idea behind prioritizing MHC binding alleles is to elicit immune response in the infected macrophages. In case of MHC-II (HTL) peptides, we relied on Tepitool and chose a pre-selected panel of 26 alleles.

### Prediction of linear B cell Epitopes

B cells also known as plasma cells or antibodies are inevitable to combat the foreign antigen and hence plays a key role in the immune cross-talk. Thus, for B-cell epitopes we used ABCpred server [14] in which B-cell epitopes were obtained from B cell epitope database (BCIPEP), which contains 2479 continuous epitopes, including 654 immunodominant, 1617 immunogenic epitopes. All the identical epitopes and non-immunogenic peptides were removed, finally we got 700 unique experimentally proved continuous B cell epitopes. The dataset covers a wide range of pathogenic group like virus, bacteria, protozoa and fungi. Final dataset consists of 700 B-cell epitopes and 700 non-epitopes or random peptides (equal length and same frequency generated from SWISS-PROT). Finally, we selected a window of 16mer peptides and a default threshold of 0.51 was chosen.

### Data cleaning and peptide Filtering

Finally, the pool of predicted peptides we obtained was scrutinized by applying several immunogenicity tests. Initially, we screened all the epitopes for their antigenicity via *VaxiJen* v2.0 server [12] (threshold of 0.4) and further determined the allergenicity profile of epitopes using AllerTop [15] which passed the antigenicity test. However, in case of MHC-I epitopes; we applied the IEDB Class-I immunogenicity (http://tools.iedb.org/immunogenicity/) test to determine if the predicted peptide was immunogenic prior to the antigenicity profile. Upon filtration of quality peptides, all the chosen peptides were finally verified for the toxic nature by using ToxinPred server [16]. Upon selection of peptides, we prioritized those epitopes which had maximum number of allele binders in order to obtain a wide population coverage across different ethnicities.

### Construction of Chimeric Vaccine

The shortlisted epitopes were assembled to construct the sub-unit vaccine design. To construct an efficient vaccine sequence; following parameters were taken into account: 1) the epitopes were 100% conserved 2) epitopes indicated no similarity with human proteins 3) a strong binding affinity was built between the peptides and respective HLA alleles 4) peptides showed maximum population coverage (for this reason, we prioritized epitopes with maximum number of allele binders as discussed earlier). Thus, we selected a suitable adjuvant which was attached to N-terminal through the EAAAK linker. The following B and T-cell epitopes were separated by flexible GPGPG linkers to formulate a tension-free reliable vaccine design [17].

### Physicochemical analysis of primary vaccine sequence

The physicochemical parameters [number of amino acids, theoretical isoelectric point (pI) molecular weight, amino acid, and atomic composition, extinction coefficients, estimated half-life, aliphatic and instability index, as well as grand average of hydropathicity (GRAVY)] of designed vaccine construct were evaluated by submitting the primary protein sequence to ProtParam web-server [18].

### Prediction of Secondary and Tertiary Structure

For secondary structure prediction, we used a combination of 2 servers namely: PSIPRED v4.0 [19] and SOPMA server [20]. Both the servers were utilized with default parameters to determine the secondary structural configurations. The 2-dimensional features included: alpha-helix, random coils, and beta-turns. The expected accuracy for SOPMA is 80% whereas, for PSIPRED v4.0, it is nearly 78%. Subsequently, the tertiary structure was predicted using SWISS-MODEL and I-TASSER.[21–23]

### Structure Refinement and Validation

The 3D structure of the chimeric vaccine was refined using the GalaxyRefine tool [24]; the server deploys an algorithm which performs thousands of repeated structural perturbations and overall structure relaxation steps through molecular simulation in order to obtain a more refined and reliable structure. The refined structure was then subjected to ProsaWeb analysis and ERRAT verification as a final passing test of structural verification.

### Molecular Docking

For molecular interaction of the proposed vaccine with the receptors, we retrieved the TLR-2 and TLR-4 structures from the Protein Data Bank (PDB) and the heteroatoms and B, C, and D chains were removed in the Discovery Studio software package. Finally, we used the ClusPro docking server to predict the confirmational binding along with lowest and center energy of the docked complexes [25].

### Molecular Dynamics Simulation

The molecular dynamics (MD) simulation of the 2 docked complexes was performed on iMOD server [26] and the outcome is presented in the results section.

### Immune Simulation

To predict the immunological response, we subjected the designed vaccine to an immune simulation performed via C-ImmSim server [27].

## Results

### Acquisition of Protein sequence and Preliminary analysis

The full-length sequence of the MPXV antigen; cell-surface binding protein comprising of 304 amino acids was retrieved from UniProt and the protein was presented with an antigenic score of 0.5311 reflecting its significant antigenic nature. Upon further analysis, the protein was confirmed to be non-allergic and non-toxic.

### Prediction of B and T-cell epitopes

Initially, a total of 26 B-cell epitopes were ranked according to their binding affinity. Upon further scrutinization through immunological tests, we filtered out 8 epitopes which were presented to be antigenic and non-allergic. The shortlisted epitopes have also been subjected to conservation analysis, hence manifesting cross protection against other species. For MHC-I epitopes, we obtained 106 epitopes initially, however, we only prioritized the epitopes having maximum number of allele binders and a significant binding score. Upon verifying the antigenicity and allergenicity, we screened 6 epitopes fulfilling the criteria. Antigenic epitopes tend to trigger a large number of antibody titers to fight the infection. Among predicted epitopes of MPX virus, five epitopes showed considerable antigenic potential, including five from the N-terminal domain and one from the C-terminal domain. For MHC-II epitopes, a total of 14 peptides were predicted initially and upon analysis we obtained 6 epitopes. Subsequently three epitopes: ^277^AIIAIVFVFILTAIL^291^, ^272^GNKTFAIIAIVFVFI^286^, ^288^TAILFLMSQRYSREK^302^ showed excellent results.

### Shortlisted epitopes for vaccine construct

The shortlisted peptides are listed in table 1.

**Table 1.**
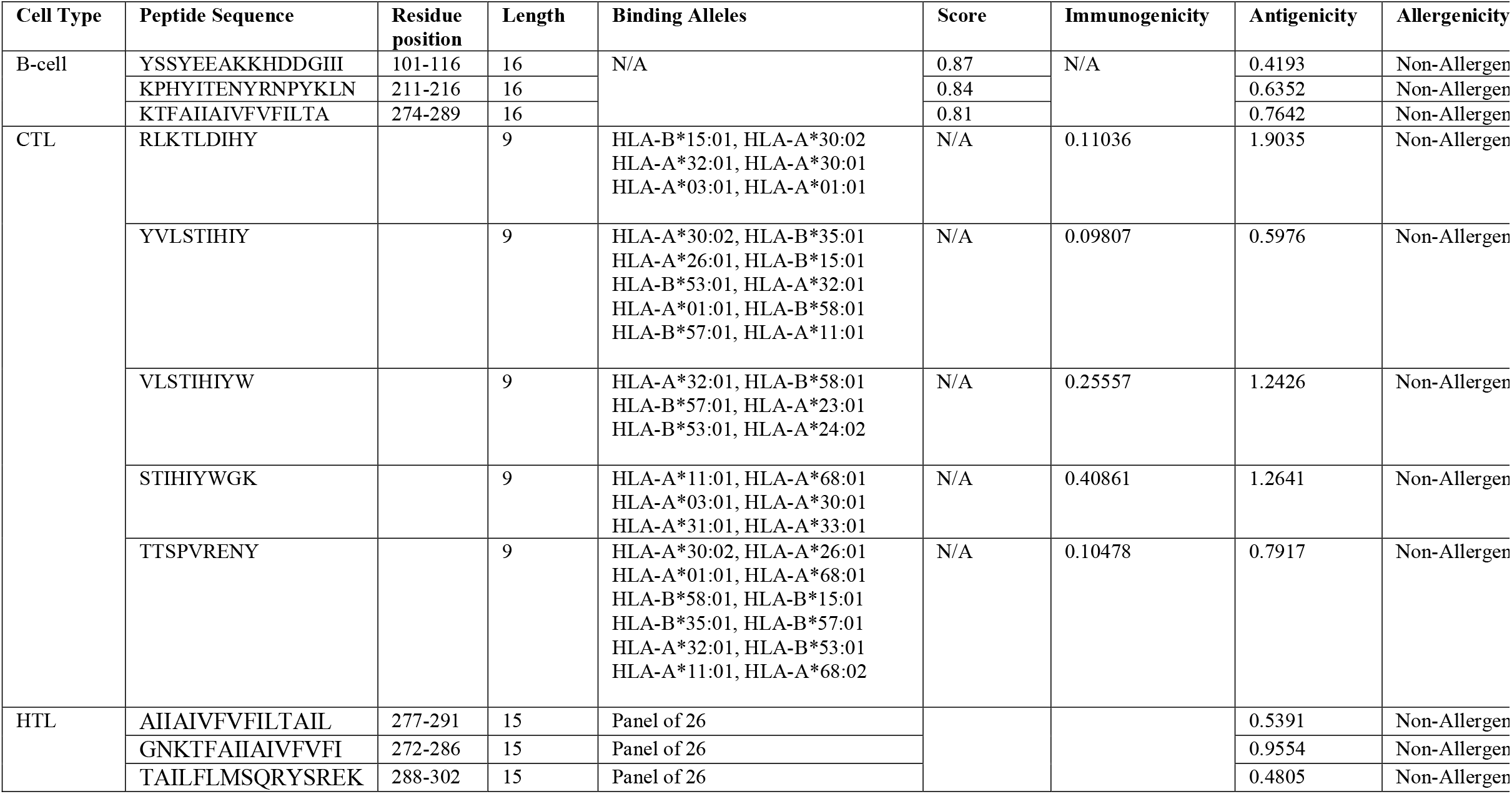
B and T-cell epitopes for final vaccine construction.

### Construction of Multiepitope sub-unit vaccine design

The final designed vaccine comprised of 337 amino acids and 11 immunogenic epitopes (3 B-cell, 5 CTL, 3HTL) shown in figure 2. The epitopes were linked together via GPGPG linkers. GPGPG linkers prevents the generation of junctional epitopes, which is a major concern in the design of multiepitope vaccines; On the other hand, they facilitate the immunization and presentation of epitopes. To prepare our vaccine peptide for a robust immune response, we attached truncated Ov-Asp1 adjuvant (IVVAVTGYNCPGGKLTALERKKIVGQNNKYRSDLINGKLKNRNGTYMPRGKNMLELTWDCKLESSAQRWANQCIFGHSPRQQREGVGENVYAYWSSVSVEGLKKTAGTDAGKSWWSKLPKLYENNPSNNMTWKVA GQGVLHFTQ) at the N-terminal through the EAAAK linker.

**Figure 2.**
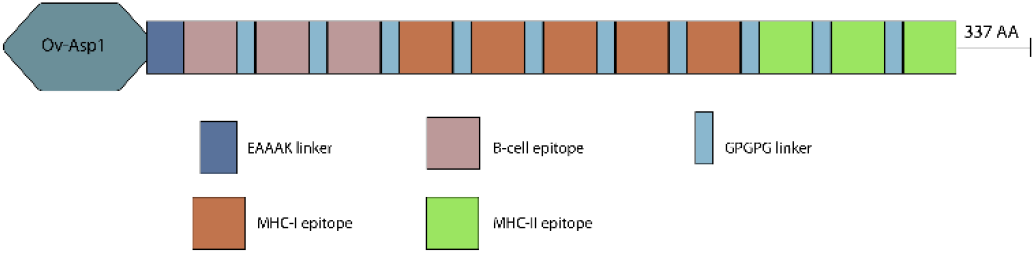
Design of chimeric vaccine. The primary vaccine map illustrates the attachment of Ov-Asp1 adjuvant to b-cell epitopes via EAAAK linker at the N-terminal. Following the b-cell epitopes, there are MHC-I and MHC-II epitopes and all the peptides are separated by the GPGPG linkers to impart overall flexibility to the final structure.

### Physicochemical Analysis of Primary vaccine design

The physicochemical parameters of the chimeric vaccine were determined through ProtParam tool on ExPasy server. Our vaccine comprised of 337 amino acid sequence with a molecular weight of 36474.94 g/mol and a theoretical pI of 9.67 which indicated the basic nature of our designed vaccine. The instability index (II) was computed to be 24.38 classifying the protein as stable. Meanwhile, the designed vaccine indicated an aliphatic index of 80.45 and a Grand average of hydropathicity (GRAVY) of −0.237 classifying our vaccine as hydrophilic in nature. In terms of half-life, the vaccine peptide indicated the estimated half-life is: 30 hours (mammalian reticulocytes, *in vitro*), >20 hours (yeast, *in vivo*), >10 hours (*Escherichia coli, in vivo*). Most importantly our vaccine protein presented to be highly soluble with a predicted scaled solubility of 0.513. Additionally, the peptide presented: 1) Total number of negatively charged residues (Asp + Glu): 19 and 2) Total number of positively charged residues (Arg + Lys): 36.

### Prediction of Secondary, Tertiary structure and 3D structure refinement, validation of chimeric vaccine

The predicted secondary structure obtained from PSIPRED (shown in figure 3) comprised 16.91% of alpha helix, 9.79% of beta turn, 37.09% of coil and 36.20% of strand. Finally, the obtained tertiary structure is shown presented in figure 4a.

**Figure 3.**
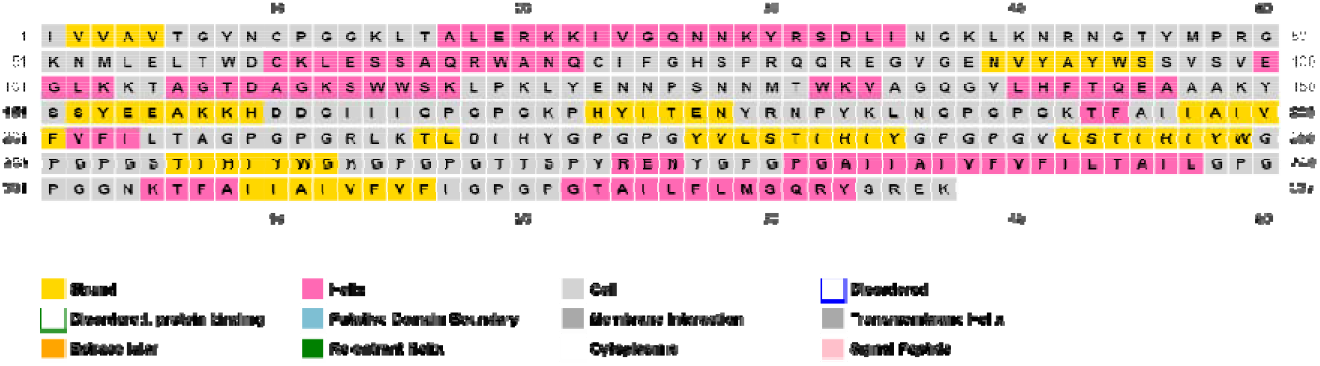
Secondary Structure of the vaccine candidate.

**Figure 4.**
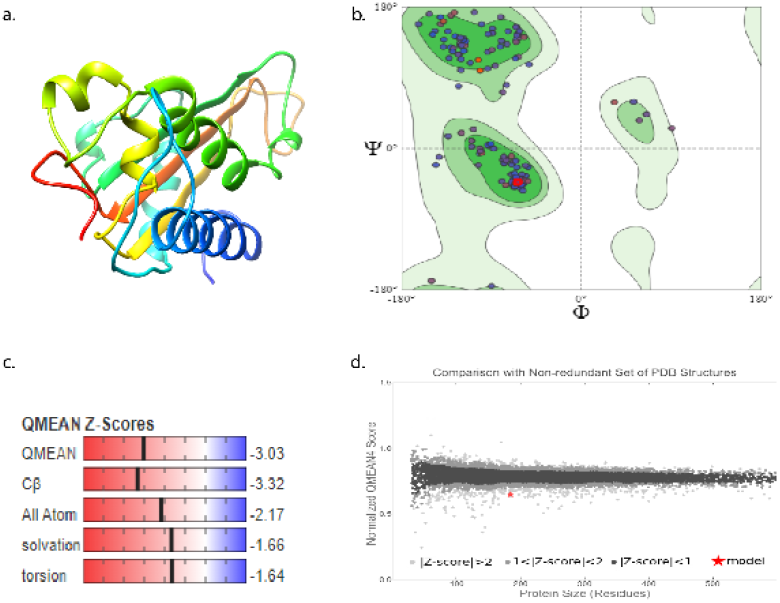
(a) 3D structure of the vaccine peptide visualized through Chimera software. (b) Ramachandran plot analysis indicated that 94.54% residues were in favored region whereas, the clash score was only 4.15% and the outliers were only 2.19%. (c) the QMEAN Z-Scores are all less than 1 indicating an overall stability of our predicted structure and (d) the scatter plot indicates that the peptide structure has a Z-score of less than 2 upon comparison with non-redundant set of a PDB structures.

The predicted 3D structure of the chimeric vaccine was refined by Galaxy Web Refinement tool and the best model indicated an enhanced percentage of 96.2 residues in favored region as compared to the previously determined 94.54%. Additionally, model 1 had a RMSD value of 0.287 and a MolProbity of 1.866. For model 1, structure perturbation is applied only to clusters of side-chains, and for model 2~5, more aggressive perturbations to secondary structure elements and loops are also applied. The ERRAT value was computed to be 82.9545 signifying the overall quality of vaccine protein and a Z-score of −3.7 from ProsaWeb (figure 5).

**Figure 5.**
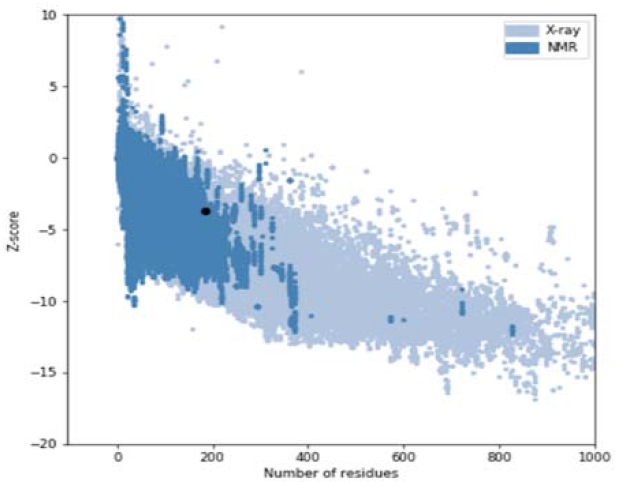
The 3D vaccine protein indicates a Z-score of −3.7 from ProsaWeb generated validation of the structure

### Molecular Docking of Proposed vaccine candidate with TLR-2 and TLR-4

The molecular interactions of the chimeric vaccine with toll like receptor-2 and 4 were assessed by the protein-protein docking performed on ClusPro server. According to ClusPro, the top cluster containing the maximum members is most reliable; however, we selected model 4 for vaccine+TLR-2 interaction because it presented with the lowest free energy binding and better structural conformation. For vaccine+TLR-4 complex, we selected model 0, as it presented with the best model scores. The center and lowest energy for both the complexes is shown in figure 6. The molecular interactions between the vaccine chain (A) and receptor chain (B) were determined by deploying the PDBsum tool. For vaccine (chain A) and TLR-2 (chain B) complex, the interface residues were 23 and 35 along with an interface area of 1757Å^2^ and 1593Å^2^ respectively. A total of 8 salt bridges and 21 hydrogen bonds were developed between the 2 proteins. The hydrogen bonds were formed between: LYS-51 and SER-29 at a distance of 2.67 Å^2^, LYS-192 and GLY-38 at 2.70 Å^2^, LYS-51 and SER-39 at 2.63 Å^2^, LYS-192 and SER-60 at 3.19 Å^2^, LYS-149 and ASN-61 at 2.72 Å^2^, ASN-186 and TYR-109 at 2.90 Å^2^, ASN-186 and ASN-130 at 3.05 Å^2^, ARG-80 and MET-159 at 2.98 Å^2^, LYS-133 and GLU-177 at 2.52 Å^2^, LYS-133 and GLU-177 at 2.61 Å^2^, TRP-132 and GLU- 178 at 2.80 Å^2^, ASN-186 and GLU-180 at 2.83 Å^2^, ARG-80 and ASP-231 at 2.83 Å^2^, PHE-75 and HIS-318 at 2.72 Å^2^, LYS-117 and GLU-369 at 2.67 Å^2^, LYS-117 and GLU-369 at 2.60 Å^2^, TYR-91 and GLN-396 at 2.60 Å^2^, TYR-93 and ASN-397 at 2.82 Å^2^, TYR-93 and HIS-398 at 2.96 Å^2^, TRP-94 and HIS-398 at 2.88 Å^2^, SER-113 and LYS-422 at 2.61 Å^2^.

**Figure 6.**
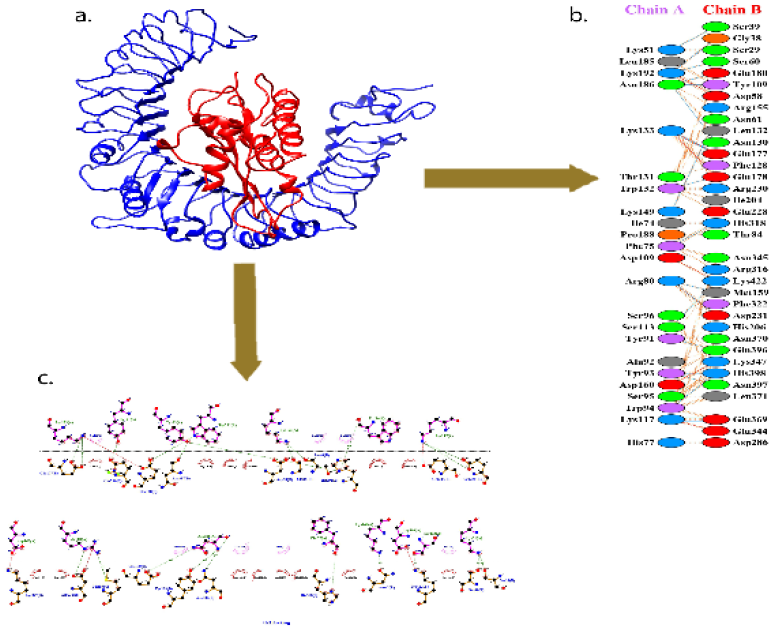
(a) Binding interaction of vaccine with toll-like receptor 2. (b) Interaction plot of vaccine (chain A) and TLR-2 (chain B). (c) Ligplot of the docked complex.

For vaccine and TLR-4 complex, there were 4 salt bridges and 27 hydrogen bonds. The hydrogen bonds developed between: LYS-38 and VAL-32 at 2.76 Å^2^, ARG-49 and GLN-39 at 2.96 Å^2^, ARG-49 and GLN-39 at 2.62 Å^2^, LYS-51 and ASP-60 at 2.60 Å^2^, LYS-51 and SER-62 at 2.53 Å^2^, GLU-84 and ARG-264 at 2.72 Å^2^, ASN-186 and ARG-289 at 2.72 Å^2^, ASN-186 and ARG-289 at 3.27 Å^2^, ASN-186 and GLU-336 at 2.97 Å^2^, TRP-132 and THR-357 at 2.84 Å^2^, LYS-184 and SER-360 at 2.92 Å^2^, ARG-80 and ASN-361 at 3.22 Å^2^, GLU-84 and LYS-362 at 2.58 Å^2^, GLN-81 and GLY-363 at 2.83 Å^2^, ARG-80 and ARG-382 at 2.62 Å^2^, HIS-77 and ASN-409 at 3.11 Å^2^, LYS-117 and GLN-430 at 2.50 Å^2^, ALA-92 and HIS-456 at 3.03 Å^2^, TRP-94 and HIS-458 at 3.13 Å^2^, SER-96 and ASN-486 at 2.84 Å^2^, TYR-91 and GLN-505 at 2.70 Å^2^, TYR-91 and GLN-505 at 3.00 Å^2^, SER-96 and GLN-507 at 2.84 Å^2^, TYR-93 and HIS-529 at 2.91 Å^2^, SER-96 and ASN-531 at 2.81 Å^2^, VAL-97 and ASN-531 at 2.94 Å^2^, SER-95 and ASN-531 at 2.92 Å^2^.

The free energy profile of docked complexes is presented in table 2.

**Table 2.**
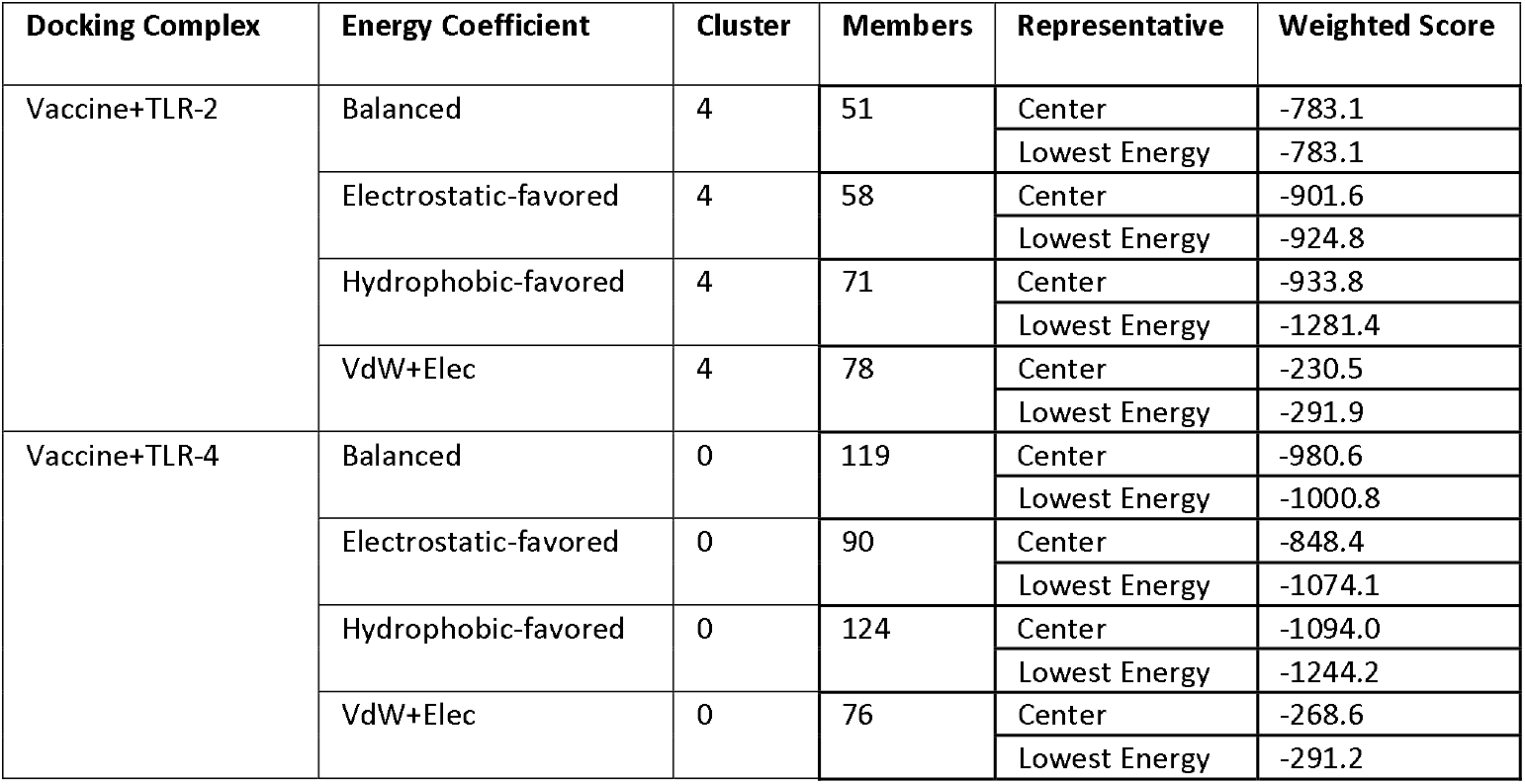
Free energy profile of the docked complexes.

### Molecular Dynamics Simulation

The results of MD simulation are presented in figure 8.

**Figure 7.**
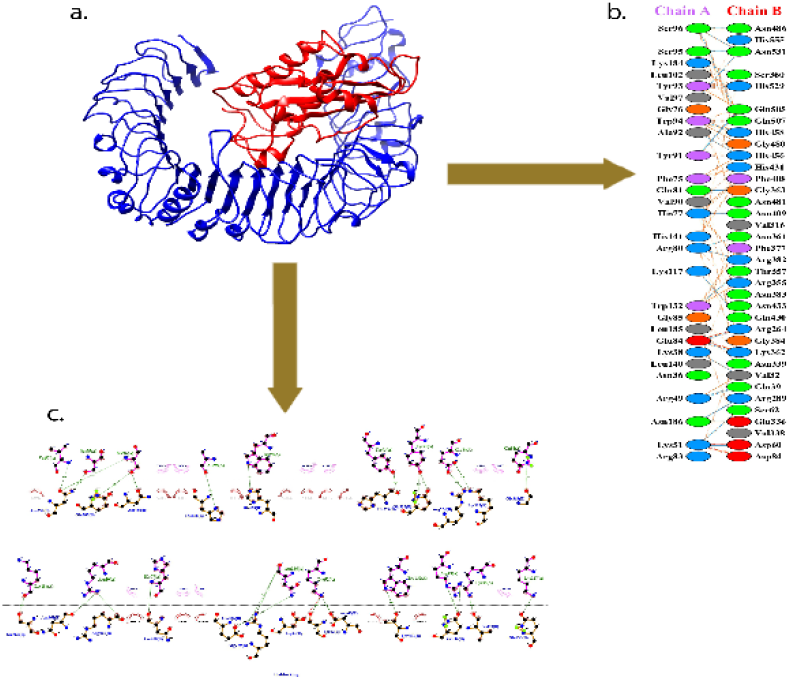
(a) Binding interaction of vaccine with toll-like receptor 4. (b) Interaction plot of vaccine (chain A) and TLR-4 (chain B). (c) Ligplot of the docked complex.

**Figure 8.**
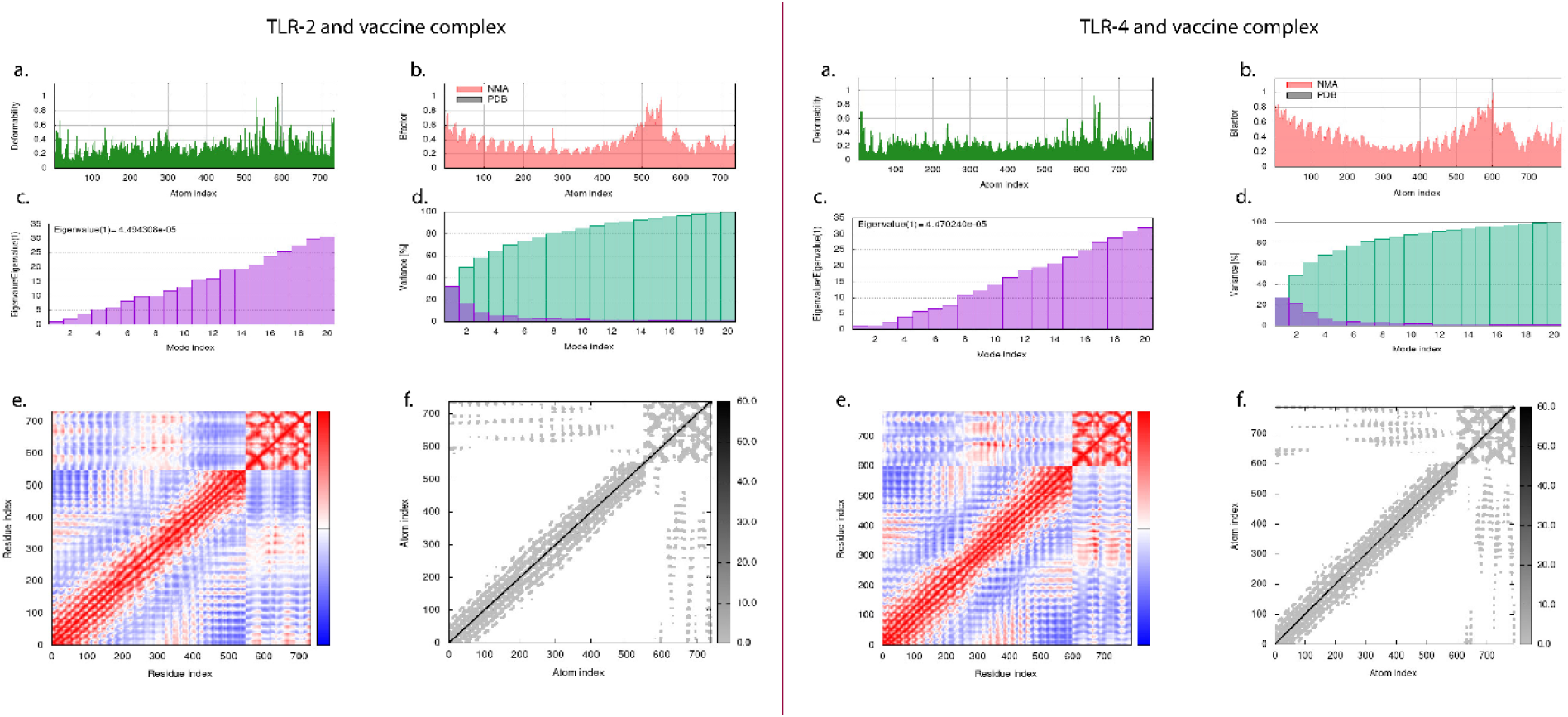
**(a) B-factor/Mobility:** The main-chain deformability is a measure of the capability of a given molecule to deform at each of its residues. The location of the chain ‘hinges’ can be derived from high deformability regions. **(b)** The experimental B-factor is taken from the corresponding PDB field and the calculated from NMA is obtained by multiplying the NMA mobility by (8pi^2———————————). Be aware that many PDB files of averaged NMR models contain no B-factors (actually, the B-factor column gives an averaged RMS). **(c) Eigenvalues:** The eigenvalue associated to each normal mode represents the motion stiffness. Its value is directly related to the energy required to deform the structure. The lower the eigenvalue, the easier the deformation. **(d) Variance:** The variance associated to each normal mode is inversely related to the eigenvalue. Colored bars show the individual (red) and cummulative (green) variances. **(e) Covariance map:** Covariance matrix indicates coupling between pairs of residues, i.e., whether they experience correlated (red), uncorrelated (white) or anti-correlated (blue) motions. **(f) Elastic network:** The elastic network model defines which pairs of atoms are connected by springs. Each dot in the graph represents one spring between the corresponding pair of atoms. Dots are colored according to their stiffness; the darker grays indicate stiffer springs and vice versa.

### Immune Simulation

The C-IMMSIM immune server was used to obtain the in silico immune profile of the given vaccination. The immune response shown to be highly significant and long-lasting. The tertiary response by the vaccine (3rd dose) was highly efficient when compared to the primary and secondary response (shown in figure). It is observed that during the 3rdshot of vaccine, the antigenic surge was already plummeted and in turn the concentration of immunoglobins (IgM+IgG, IgM, IgG1+IgG2, IgG1, IgG2) indicated a significant increase. Alongside, various isotypes of the B-cells were found (isotype IgM, IgG1, IgG2) indicating the isotype switching and the memory B-cells were found to reach the maximum potential after the 3rd dose of the vaccine. In addition, our vaccine propounded the maximum number of active B-cell population and highly low concentration of anergic B-cells were indicated which reflects the immunogenic nature of our vaccine. Similarly, there was a significantly higher Th and Tc cell response right after the complete course of vaccine shots. A similar trend was also observed in the population of macrophages, natural killer cells and dendritic cells.

**Figure 9.**
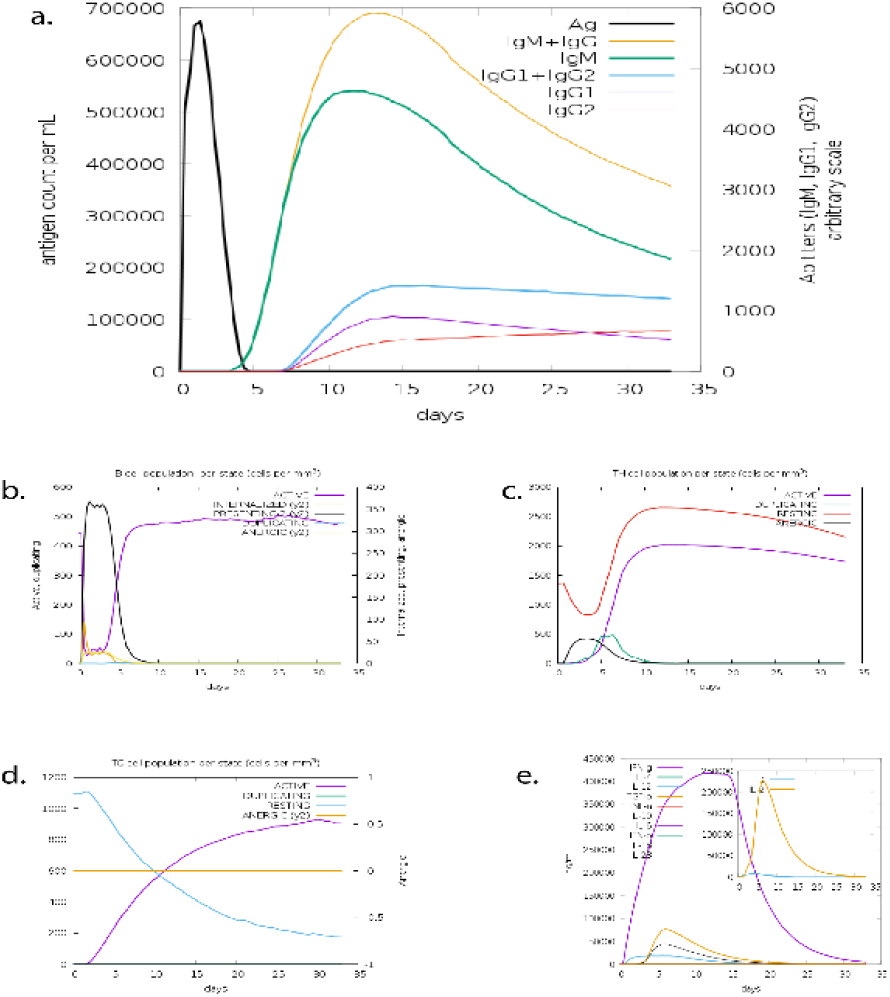
Results of Immune Simulation. (a) Immunoglobin profile indicates a significant rise in the IgM+IgG titer after 5 days. (b) The B-cell concentration in the active phase continuously rises after the vaccine administration followed by (c) CD4 T-helper lymphocytes in active phase and its count sub-divided per entity-state (i.e., active, resting, anergic and duplicating). (d) Meanwhile, the CD8 count shows a rising concentration in the active phase. (e) Cytokines: Concentration of cytokines and interleukins. D in the inset plot is danger signal.

## Discussion

The monkeypox virus cases recorded in May 2022, suddenly became a public health threat after the emergency by World Health Organization. Due to the destruction by Covid-19 pandemic, researchers and health experts have become alert in order to combat this disease to avoid the potential pathogen of transpiring as a next global health issue [28]. However, being a member of orthopoxviridae family, the pathogen is less severe than smallpox but implicates serious complications in immunocompromised individuals. Although fatalities by monkeypox are rare with only 22 deaths out of more than 57000 confirmed cases of virus, but it should not be neglected in the first place and effective treatments and therapeutics should be a priority [29]. In terms of prevention, data has suggested that prophylactic vaccines licensed for smallpox may be effective against smallpox by improving the clinical manifestations of the infection. Currently, there are 2 specific vaccines and 1 suggested upon investigational new drug (IND) protocol is available for the smallpox disease [30–32]. Out of these, JYNNEOS™ (a live viral vaccine developed through the modification of vaccinia Ankara-Bavarian Nordic; MVA-BN strain) is considered an effective alternative vaccine therapy for smallpox. Notable, the previous data has suggested that this vaccine has been 85% effective against the monkeypox virus [30, 33]. Regardless of the existing alternative prior immunization therapies, there needs to be a specific treatment for the infection considering the highly mutating nature of the viruses.

Thus, the current study has employed reverse vaccinology approach using the immunoinformatics-guided principles to design a chimeric vaccine against the monkeypox virus. This approach is highly acceptable in the present era to avoid extra costs, resources and formulate effective target-specific therapeutics in a timely manner. A much interest has been developed and immunoinformatics-guided techniques has been used to formulate epitope-based subunit vaccines [34–36].

Aiming to design a suitable vaccine against monkeypox, we deployed several computational tools. For preparation and processing of virus antigen as a target, we retrieved the amino acid sequence of cell surface-binding protein from UniProt. Cell surface-binding protein is responsible of vital biological processes for MPXV such as viral entry into host cell, virion attachment to host cell and host-virus interaction [37]. Thus, the present study aimed to design a multi-epitope subunit vaccine that can elicit an immune response against MPXV using cell surface-binding protein as a target. Since B-cell lymphocytes are involved in the humoral immune response against the target antigen, so we employed the ABCpred server to retrieve 16mer B-cell epitopes and the peptides were shown to be significantly antigenic and non-allergen when passed through the computational prediction by *Vaxijen* and AllerTop server respectively. Subsequently, for a broad range coverage of peptides, we chose the Tepitool from IEDB to retrieve MHC class-I epitopes for a panel of 27 most frequent alleles. Upon further securitization, we prioritized epitopes that comprised of a higher number of binding alleles in order to cover a wide range of population. On the similar pattern, we analyzed the predicted MHC class-II epitopes from Tepitool. These epitopes were linked together by GPGPG linkers to confer better flexibility to the overall structure and finally adjuvanted with Ov-Asp1 (linked through the attachment of EAAAK linkers) for a robust immune response. The post vaccine construct analysis including physicochemical properties (on ProtParam) and solubility profile (ProteinSol) presented excellent results of the designed chimeric vaccine.

The secondary structure of the design protein sequence retrieved from PSIPRED was analyzed on SOPMA server which indicated the presence of alpha helix, beta turns and random coiled structures. Upon conformation of primary analysis, we employed the SWISS-MODEL server for prediction of 3D structure of our vaccine peptide. According to the predicted 3D structure, our vaccine had higher proportion of residues in the favorable region (Ramachandran plot) and it was further refined by the Galaxy web refinement tool for a better threaded and modified structure of the protein. The interactions between antigenic molecules and immune receptor molecules are critical for efficient antigenic transport and immune response activation. Finally, the validated structure was docked with TLR-2 and TLR-4 receptors to analyze the types of interactions between the 2 proteins. The PDBsum analysis of our docked complexes indicated that the TLR-2 had 21 hydrogen bonds with the vaccine protein whereas, the TLR-4 developed 27 hydrogen bonds. At last, the structural motion of docked complexes was analyzed by subjecting the structures to MD simulation.

## Conclusion

The present study has aimed to provide a design of highly putative vaccine candidate using the reverse vaccinology approach. The designed vaccine based on the rationale target protein is proven to be highly effective contrary to traditional vaccines with several drawbacks including toxicity and allergic reactions with lower efficacy of the administered vaccine. The peptides selected for our present multiepitope vaccine are highly immunogenic, antigenic and non-allergens; thus, provoking a robust immune response against the target cell surface-binding protein. The designed study, will be highly applicable to the future formulation of any prophylactic or therapeutic therapies against the monkeypox virus disease.

## Acknowledgement

We would like to acknowledge ViralZone (https://viralzone.expasy.org/) for **Figure 1**.

